# Not dead yet: Diatom resting spores can survive in nature for several millennia

**DOI:** 10.1101/285122

**Authors:** Anushree Sanyal, Josefine Larsson, Falkje van Wirdum, Thomas Andrén, Matthias Moros, Mikael Lönn, Elinor Andrén

## Abstract

We show for the first time the revival, viability and germination rate of resting spores of the diatom *Chaetoceros* deposited in sub-seafloor sediments from three ages (recent: 0-80 years; ancient: ∼1300 and ∼7200 calendar year before present. Sanger sequences of nuclear and chloroplast markers were performed. Our findings showed that ∼7200 calendar year BP old *Chaetoceros* resting spores are still viable and the physiological response pertaining to vegetative reproduction in recent and ancient resting spores vary. The time taken to germinate is three hours to 2-3 days in both recent and ancient spores but the germination rate (%) of the ancient spores decrease with increasing time. Based on the morphology of the germinated vegetative cell we were able to identify the species as *Chaetoceros muelleri*.

Studies of revived resting spores of marine diatoms will serve as excellent proxies of environmental change in marine environments and enable us to reconstruct ∼7000 years of diatom evolution in relation to changes of their environment. Comparison of resurrected populations obtained from these natural archives of diatoms can provide predictive models to forecast evolutionary responses of populations to environmental perturbations from natural and anthropogenic stressors, including climate change over longer timescales.

## Introduction

Human domination of the Earth’s ecosystems has accelerated in recent decades, to the extent that this period has been termed as the Anthropocene (Waters et al. 2016). Phytoplankton forms the basis of the marine food web and hence if we understand how phytoplankton will respond to environmental and climate change we will have a better chance to understand ecosystem change. Changes in species abundance with increased numbers of planktic, resistant, toxic, and introduced species due to nutrient enrichment and resulting symptoms of eutrophication, hypoxia, metal pollution and acidification (Yasuhara et al. 2012) has resulted in significant loss of biodiversity in the marine environment. We have very little knowledge about how the Earth’s biota will be affected both directly (Hofman et al. 2015) and evolutionary (Larsson et al. 2016; Whitehead 2014) by human-induced climate change, increased pollution and eutrophication. Understanding how the global biodiversity and their adaptive responses will be affected by anthropogenic perturbations has emerged as a major challenge to humanity (Hofman et al. 2015; Schlüter et al. 2014). Our current inability to do so hampers our understanding of how the future ocean will function.

Resurrection ecologists have long recognized sediments as sources of viable propagules (“seed or egg or resting spore banks”) for studying ecological and evolutionary responses (Cáceres and Hairston 1998). Resting stages in the sediment are of ecological and paleoecological importance as they can be revived when exposed to suitable environmental conditions and used as a source of genetic material for microevolutionary studies (Ellegaard and Ribeiro 2017). Studies predicting how evolution will shape the genetic architecture of populations coping with present and future environmental challenges has primarily relied on investigations through space, in lieu of time. Yet, dormant propagules in sediments are natural archives from which adaptive trajectories of populations could be traced along extended time periods. Currently, only a century old diatom resting spores have been revived (Härnström et al. 2011), and studies on dinoflagellates have shown that the survival periods of several marine dinoflagellate resting stages ranging from several months to 100 years (Miyazono et al. 2012). Previous studies on *Daphnia* and *Sphagnum* resurrected ∼ 700 years old ancient eggs (ephippia) and spores (Yousey et al. 2018, Bu et al. 2017). Hence, we lack a model system to study the changes occurring over longer evolutionary timescales. A key to understanding the adaptive capacities of species over evolutionary time lies in examining the recent and millennia old resting spores buried in the sediments.

Resting spore formation in diatoms is an effective strategy to survive periods of stress and has enabled diatoms to withstand events of mass extinction during the end of Cretaceous period (Kitchell et al., 1986). Climate, nutrient concentration, anthropogenic disturbances and physical oceanographic conditions can influence the distribution, composition and abundance of diatom species (Crosta et al. 1997) and render them a useful probe to study environment induced adaptations over long timescales (Orsini et al. 2013; Burge *et al*. 2017). Furthermore, DNA sequence data obtained from these natural archives of resurrected organisms, combined with the next generation DNA sequencing methods and models for analyzing both ecological and population genomic data can help predict the evolutionary responses of natural populations to environmental changes resulting from natural and anthropogenic stressors, including climate change (Orsini et al. 2013).

Globally, *Chaetoceros* is an abundant and diverse marine planktic diatom genus which plays a major role in marine primary production (Malviya et al. 2016). Studies in the Baltic Sea Area show that the present distribution of *Chaetoceros* species correlates with the salinity gradient resulting in higher diversity in the more marine Kattegatt and Danish straits and lower diversity in the brackish central Baltic Proper and in the nearly freshwater Bothnian Bay (Andersen et al. 2017; Fig. 1). Due to its dissolution-resistance and a high sedimentation rate (which would result in quick burial of the spores) in the Baltic basin, *Chaetoceros* resting spores are found throughout the sediment stratigraphy from when marine water entered the Baltic proper (named the Littorina Sea, ∼7500 years ago) till date (Andrén et al. 2000; Witak et al., 2011).

**Fig. 1.**
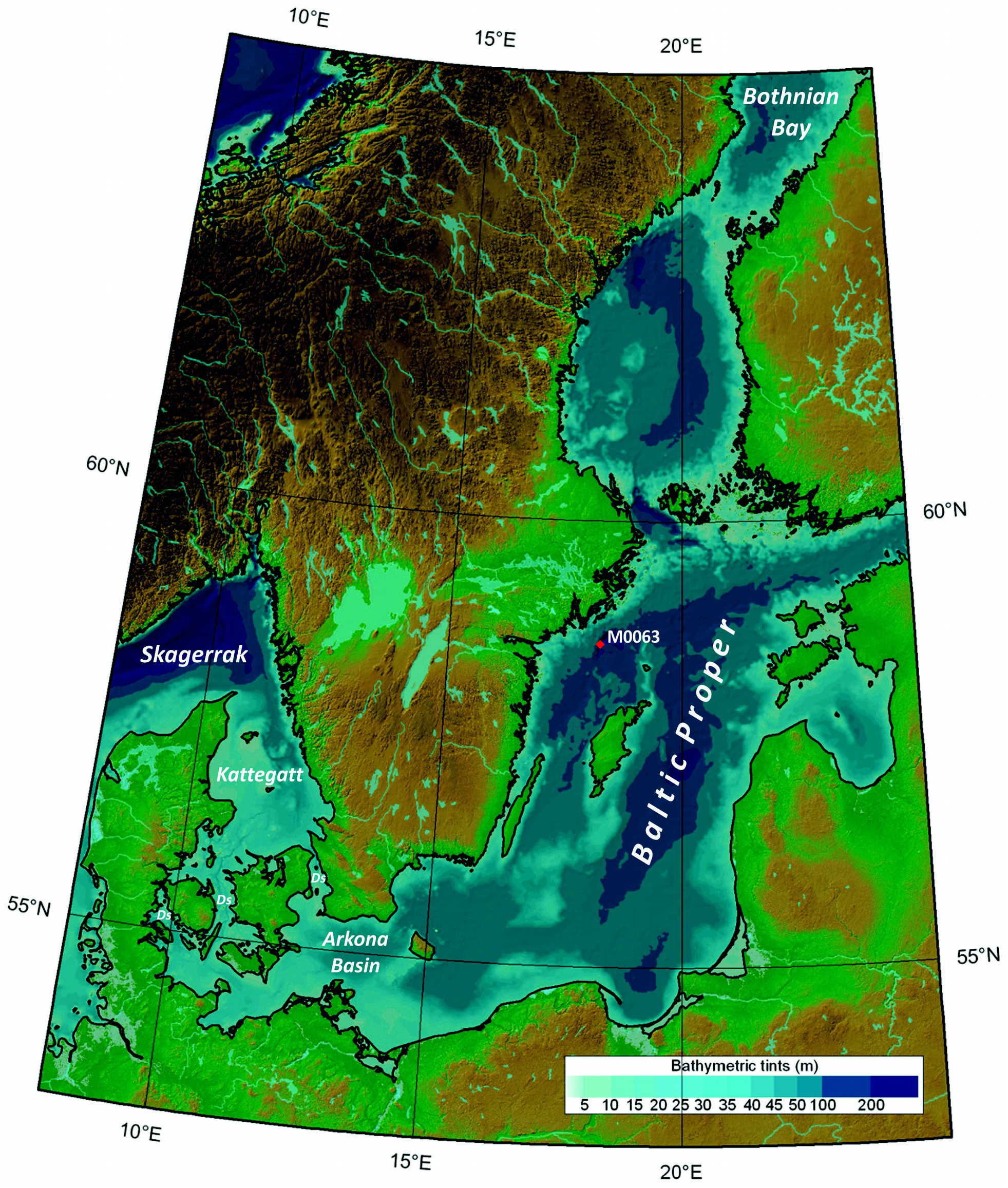
Map showing the position of the investigated IODP Expedition 347 site M0063 in the Landsort Deep (58°37.32’N, 18°15.24’E) where a ca 116 m long sediment core was drilled at a water depth of 437 m (Andrén et al. 2015). Baltic Sea has a long salinity gradient in the surface waters ranging from ∼12-30 in Kattegatt, ∼10-23 in Danish straits (Ds), ∼8-11 in Arkona Basin, 5-7.5 in Baltic Proper and ∼2-4 in Bothnian Bay (Andersen et al. 2017).

The Baltic Sea ecosystem has strong horizontal (salinity, temperature) and vertical (also oxygen) environmental gradients (Snoeijs-Leijonmalm and Andrén, 2017). The vertical salinity gradient results in stratification which together with ongoing eutrophication causes large areas with hypoxic and anoxic (oxygen concentration <2 and <0 mg L-1) bottom waters (Carstensen et al. 2014). The Littorina Sea (∼7500-3000 cal yr before present (BP)) reached a maximum surface water salinity of 12-13 in the Baltic Proper, compared to present day 5-7.5 (Gustafsson and Westman 2002). The higher salinity coincides with the Holocene Thermal Maximum (HTM), with warm and dry climate dated to ∼8000-5000 cal yr BP (Seppä et al. 2009). Also, post-deglaciation of Scandinavia, interactions between global sea level rise and isostatic adjustments resulted in different freshwater and brackish water stages, during a geologically and evolutionarily short period of time (Andrén et al. 2011; Snoeijs-Leijonmalm and Andrén, 2017).

In the open Baltic Proper, high *Chaetoceros* resting spore abundances are found in the laminated, hypoxic sediments deposited during three time periods: the last century, the Medieval Climate Anomaly (MCA, ∼1000-700 cal yr BP, Mann et al. 2009) and the HTM (Andrén et al. 2000). The abundances of resting spores in sediments from the Baltic Basin are attributed to high primary production and eutrophication (Andrén et al. 1999; 2000) and also coincides with high temperatures and higher surface water salinities (Zillén et al. 2008), at least during the HTM.

To systematically study the effects of anthropogenic perturbations over long time scales we need (1) a model organism whose resting spores have accumulated over a long period of time and (2) its ecosystem where the deposited resting spores have been subjected to anthropogenic perturbations over long timescales. The overall objective of the present study is to determine if *Chaetoceros* resting spores preserved in Baltic Sea sediments is a good model system to study the impact of anthropogenic perturbations over long time scales. The specific aims are to examine if the resting spores from three ages (recent, MCA and HTM) can be (i) revived by germinating them (ii) DNA extracted, amplified and sequenced, and (iii) identified to species level (since many spores have identical morphology in comparison to germinated living cells). In this study we identify and present a unique model system, *Chaetoceros* resting spores from the Baltic Sea, which will help understand the effects of anthropogenic alterations on the evolutionary fate of natural populations on an evolutionarily significant temporal scale instead of spatial scale.

## Materials and Methods

### Sampling

The Integrated Ocean Drilling Program (IODP) Expedition 347 drilled the Baltic Proper from R/V *Greatship Manisha* during September to November 2013. The M0063 site (58°37.32’N, 18°15.24’E) in the Landsort Deep, the deepest part of the Baltic Proper at a water depth of 437 m was drilled during this expedition (Fig. 1). Five holes (M0063A to M0063E) were drilled, using an advanced piston corer with perfluorocarbon added in the liner fluid to trace possible contamination, down to a diamicton (till) was reached at about 92 meters composite depth (mcd) (Andrén et al. 2015). The sediments were stored in the dark in a cold room at 4°C. All Expedition 347 cores were split and sub-sampled during the onshore science party at IODP Core Repository in Bremen, Germany, January to February 2014. Sediment samples used for revival experiments were requested from holes C and D in March 2015 and sub-sampled by the curators at IODP Core Repository in Bremen. The drilling site was revisited in June of 2014 with M/S *Fyrbyggaren* and the topmost unconsolidated sediments were sampled with a 1-m gravity corer. Two cores 87 cm (named M0063H, N58°37.34’, E18°15.29’) and 63 cm (named M0063I, N58°37.35’, E18°15.25’) respectively were retrieved and immediately subsampled in 1-cm slices and stored in the dark in a cold room at 4°C.

### Lithology and sample selection

The lithology at site M0063 is divided into seven lithostratigraphic units very briefly described as; below 92 mcd diamicton, 92 – 32 mcd laminated clays on cm-scale which gradually change to mixed clays in the upper part, 32 – 26 mcd homogeneous clays with sulphide banding, 26 – 0 mcd organic rich clays with more or less pronounced laminaes on mm-scale (Andrén et al. 2015).

Based on the results from the onshore science party (Andrén et al. 2015), samples from three sediment sections of different age with anticipated high spore abundance were selected to trace the genetic diversity and the evolutionary changes in the *Chaetoceros* populations: the last century (M0063I 0-63 cm), MCA (M0063C 5.42-6.32 mcd) and the HTM (M0063D 21.63-26.05 mcd).

### Dating and age modelling

The two old selected time intervals corresponding to the MCA (M0063C) and the HTM (M0063D) were dated by radiocarbon accelerator mass spectrometry (AMS) at Beta Analytic, USA.

In order to assign a calendar year age to each level analyzed an age-depth modeling was performed using the software CLAM version 2.2 (Blaauw, 2010) with 2000 iterations and a custom-built calibration curve based on the *IntCal13* calibration dataset (Reimer et al. 2013) with a mean and standard deviation of 900 and 500 ^14^C years respectively (Obrochta et al. 2017). The sediment surface is assumed to be modern (i.e. 2013, the year of the coring).

The youngest sediment (the two short gravity cores M0063H and M0063I), was dated using stratigraphic time markers (Moros et al. 2017). In order to identify stratigraphic time markers, mercury (Hg) and artificial radionuclide (^137^Cs and ^241^Am) measurements on M0063H have been performed. Core correlation of M0063H and M0063I is based on characteristic features of the mercury downcore profiles. Hg and ^137^Cs were normalized to bulk organic carbon (TOC) in order to eliminate the dilution effect of the massively occurring manganese-carbonate layers.

### Resting spore concentration

To calculate the concentration of resting spores in the upper 30 m (mcd) from hole M0063D, sediments were freeze-dried, and a known weight of sediment (on average ∼0.1 g) was subsampled. Cleaning of diatoms were performed according to standard procedures (Battarbee 1986). Microspheres were added in the last step to allow for the calculation of resting spore concentrations before mounting for permanent slides in Naphrax™ (Battarbee and Kneen 1982). A total of 54 samples were analyzed for diatom spores using light microscopy at 1000x magnification and immersion oil. Due to the difficulties in differentiating between the morphology of *Chaetoceros* resting spores, they were not separated to species level. Concentrations of *Chaetoceros* resting spores were calculated and expressed in numbers of valves per gram dry weight (gdw) (Battarbee and Kneen, 1982).

### Single resting spore isolation from several millenia old resting spores

Resting spores were isolated from recent sediment samples (∼the last century) as well as from MCA and HTM ages. The isolation procedure was modified from Throndsen (1978) were each sample was diluted with ∼1ml of sterile ddH_2_O in a 30mm × 15mm petri dish and a drop of the solution was placed on a concave microscope slide. Individual resting spores were isolated through manual suction using 20–40 μl drawn-out disposable pipettes and examined on a concave microscopic slide under an inverted microscope with a 10x objective and a WF40x eyepiece with a total magnification of 400x. The isolated resting spore possibly with associated contaminants was transferred to a new ddH_2_O water droplet. This isolation and transfer was repeated 2–5 times to remove any contaminants. Individual resting spores were then isolated for the final time and transferred to a 30mm × 15mm petri dish containing artificial seawater medium (Tropic marine^®^ Wartenberg, Germany).

### Germination and germination rate of the recent and several millenia old resting spores

The resting spores from the last century, MCA and HTM ages were germinated in 30mm × 15mm petri dishes containing artificial seawater medium with salinities ranging from ∼10.5 to ∼11 at temperatures ranging between 15 to 18°C.

The germination rates of the resting spores were estimated by counting the number of germinated spores that germinated in a field of the microscope. The germination rate observed in 12 to 21 microscopic fields were recorded for each sample at a total magnification of 400X. The average germination rate for the recent and ancient resting spores were calculated. The germination rate was analyzed using a one-way ANOVA with the age of the resting spores as the fixed effect and the germination rate as the dependent variable using R (R Core Team, 2017).

### Growing the unialgal Chaetoceros cultures

Unialgal cultures of *Chaetoceros* from revived individual resting spores from the last century were established from an inoculum from the petri dish which was transferred to a 50 ml angled neck Nunc^®^ EasY Flasks^™^ with vent caps. The flasks were filled with the Guillard’s (F/2) Marine Water Enrichment Solution which contained silicate and grown in growth chambers with controlled temperature (15 −18°C) and a 12:12 light-dark regime. The culture media was prepared by adding 20 ml of the Guillard’s F/2 medium in 1L of artificial seawater. The light intensity was 125 *u*E m^−2^ s^−l^ and was provided by fluorescent lamps. The cultures were subcultured every two weeks to maintain the culture.

### Identification of the Chaetoceros species

The germinated *Chaetoceros* species was cleaned in hydrogen peroxide according to the method described in Battarbee (1986), dried onto a coverslip and mounted in Naphrax™ (refraction index n_D_=1.73). Qualitative analysis was carried out with an Olympus BX51 light microscope using Nomarski differential interference contrast with a magnification of 1000x and oil immersion. Diatom species identification followed Cleve-Euler (1951), Ishii et al. (2011), Johansen and Rushforth (1985) and Krammer and Lange-Bertalot (1991).

### DNA isolation from diatom cultures of recent resting spores

Total DNA was extracted from each *Chaetoceros* culture (based on revived spores from the last century) as described below. 25 ml of the cell cultures were centrifuged at 2500 rpm for 10 min in a 50 ml Falcon tube, 20 ml of the supernatant was discarded using a syringe. The remaining 5 ml were centrifuged again at 2500 rpm for 5 min. 4 ml of the supernatant was discarded. The remaining 1 ml was spun for 2 minutes. The supernatant was discarded. Genomic DNA was extracted using a modified chloroform method with the addition of cetyltrimethylammonium bromide (CTAB) as describe by Zuccarello and Lokhorst (2005). The culture pellet was grinded with microfuge pestle in 500 μl of CTAB extraction buffer (2% CTAB, 0.1 Tris-HCl (pH 8.0), 1.4 M NaCl, 20 mM EDTA, 1% PVP), 2 μl RNAse A (100mg/ml) and 5 μl Proteinase K (20 mg/ml) and the tubes were incubated at 55-60°C for 30 min. Equal volume of chloroform:isoamyl alcohol (24:1) was added and mixed. The tubes were then spun at 12,000 g for 5-10 min. The supernatant was removed to a new tube, avoiding interface. This step was repeated. DNA was precipitated in 100% ice-cold isopropanol, each tube was inverted and placed at room temperature for 30 min. The tubes were spun for 20 min at 12,000 g. The DNA pellet was washed in 500 μl of 70% ethanol and the tubes were spun for 5 min. The supernatant was poured out and the DNA pellet was air-dried. The DNA pellet was suspended in 50 μl of 0.1 X (0.1mM EDTA, 1 mM Tris) TE buffer. The quality of the DNA extraction was assessed by visualizing the products on a 1.5% agarose gel and the DNA concentration was evaluated with a nanodrop. The DNA was then frozen at −80°C for subsequent PCR and sequencing reactions.

### DNA isolation from single millenia old Chaetoceros resting spores

Single cell Chelex® DNA extraction was performed on spores which were obtained from the sediments of MCA age by using the Chelex method. For DNA extraction, Chelex-stored samples were incubated for 20 min at 95°C. They were then vortexed for 15 s and centrifuged for 15 s at 12,000 g. DNA extraction from the germinated resting spores of the HTM age will be done in a future study.

### PCR and sequencing

For the PCR reaction, 25 μL of reaction mixture contained as a final concentration, the two primers at 15 pmole each; 5–20 ng DNA template; 0.2 mM each dATP, dCTP, dGTP and dTTP; 2.5 μL of 10×PCR buffer for Blend Taq (25 mM MgCl_2_ concentration); and 0.1 units of Blend Taq polymerase (Thermo Scientific). The PCR condition was as follows: an initial step at 94°C for 2 min followed by 30 cycles with a denaturation temperature of 94°C for 30 s, an annealing temperature of 53°C for 30 s and an extension temperature of 72°C for 1.5 min. The PCR was assessed by visualizing the products on 1.5% agarose gel. Six primer pairs from nuclear ribosomal RNA (SSU) and chloroplast DNA (*rbcL*) from three unialgal cultures were amplified (Table 3). Sanger sequencing was performed on the PCR products (Macrogen inc.) of six nuclear ribosomal RNA (small (SSU) and chloroplast DNA (*rbcL*) markers which was used to amplify *Chaetoceros* species (Lee et al. 2013) (Table 3).

### Analysis of DNA sequences

The sequences were blasted using the nucleotide BLAST tool of NCBI. Sequences were aligned by using Molecular Evolutionary Genetics Analysis version 5 (MEGA5) (Table 3).

## Results

### Dating and age modelling

Two sediment samples of MCA age and four samples of HTM age were radiocarbon dated (Table 1) and used for age modelling to assign an age to all individual samples used for the resurrection study (Table 2). Radiocarbon dating together with stratigraphic time markers for the youngest sediment (Fig. 3) showed that our samples were from three distinct time intervals from present time (0 years) to 7200 cal yr BP. The sediment from the short gravity core M0063I with an anticipated age of the last century could be assigned an age of ca. 1940-1935 (i.e. ∼0 - 80 years) at 63 cm, based on the stratigraphic time markers identified (Fig. 3). The sediment samples used for the resurrection study gave after age modelling ages of ∼1300-1000 cal yr BP from MCA age (M0063C 5.42-6.32 mcd) and 7200-6000 cal yr BP of HTM age (M0063D 21.63-26.05 mcd) (Table 2). Age modelling made it possible to estimate the calendar years age for all levels analyzed except for the sample at 6.32 mcd which was outside the dated interval. A linear extrapolation from the sample at 6.22 mcd was used to estimate the age of this sample (Table 2).

**Fig 2.**
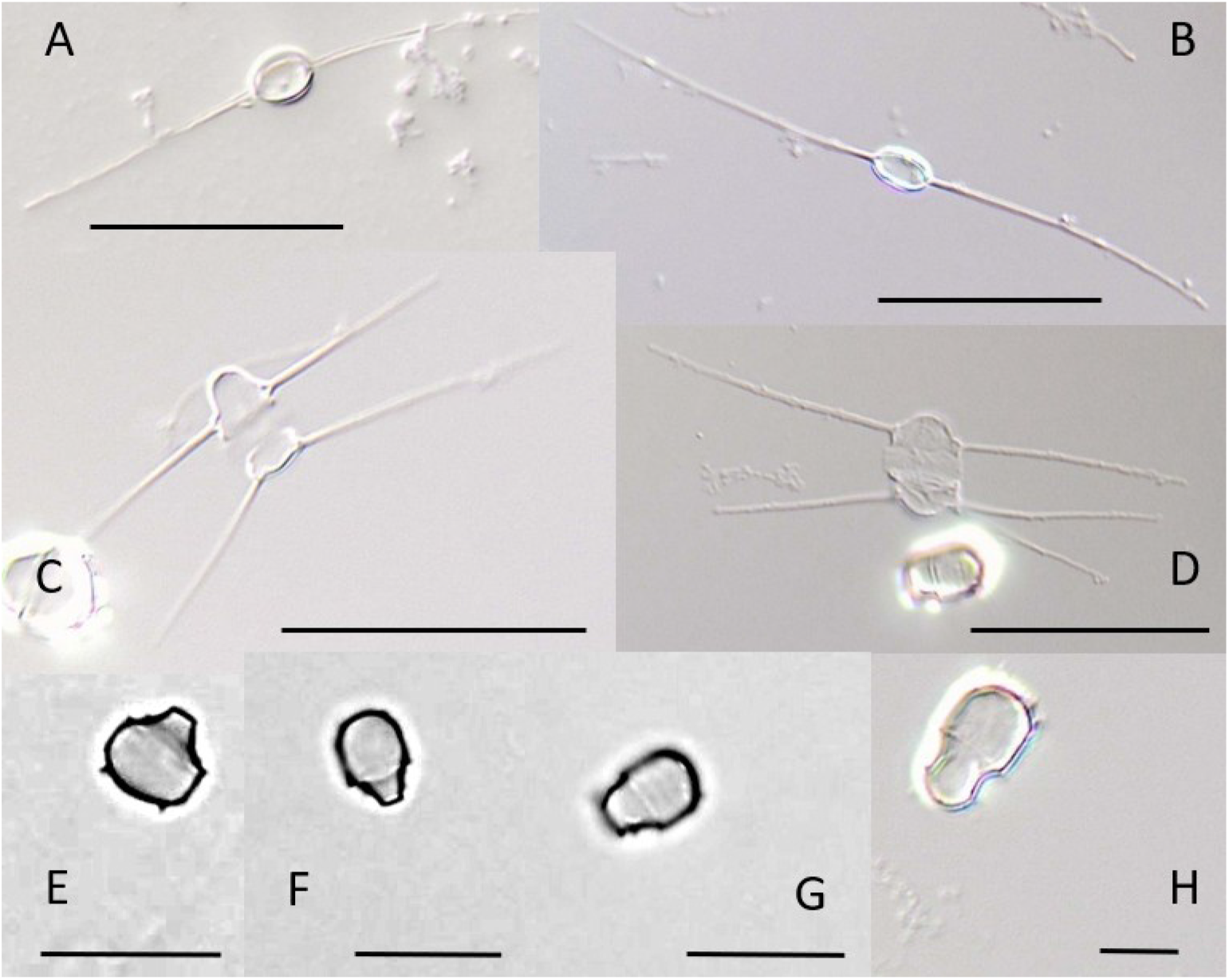
Light micrographs of *Chaetoceros muelleri* Lemmermann in the culture revived from the sediment at 10-11 cm depth. A-D. Vegetative cells in valve view (A-B Smooth solitary cell) and girdle view (C-D showing convex valve shape). Note that both vegetative cell and resting spore is visible in D. E-H. Resting spores in girdle view showing various characteristic morphologies. Scale bar A-D 20 microns, E-H 5 microns.

**Fig. 3.**
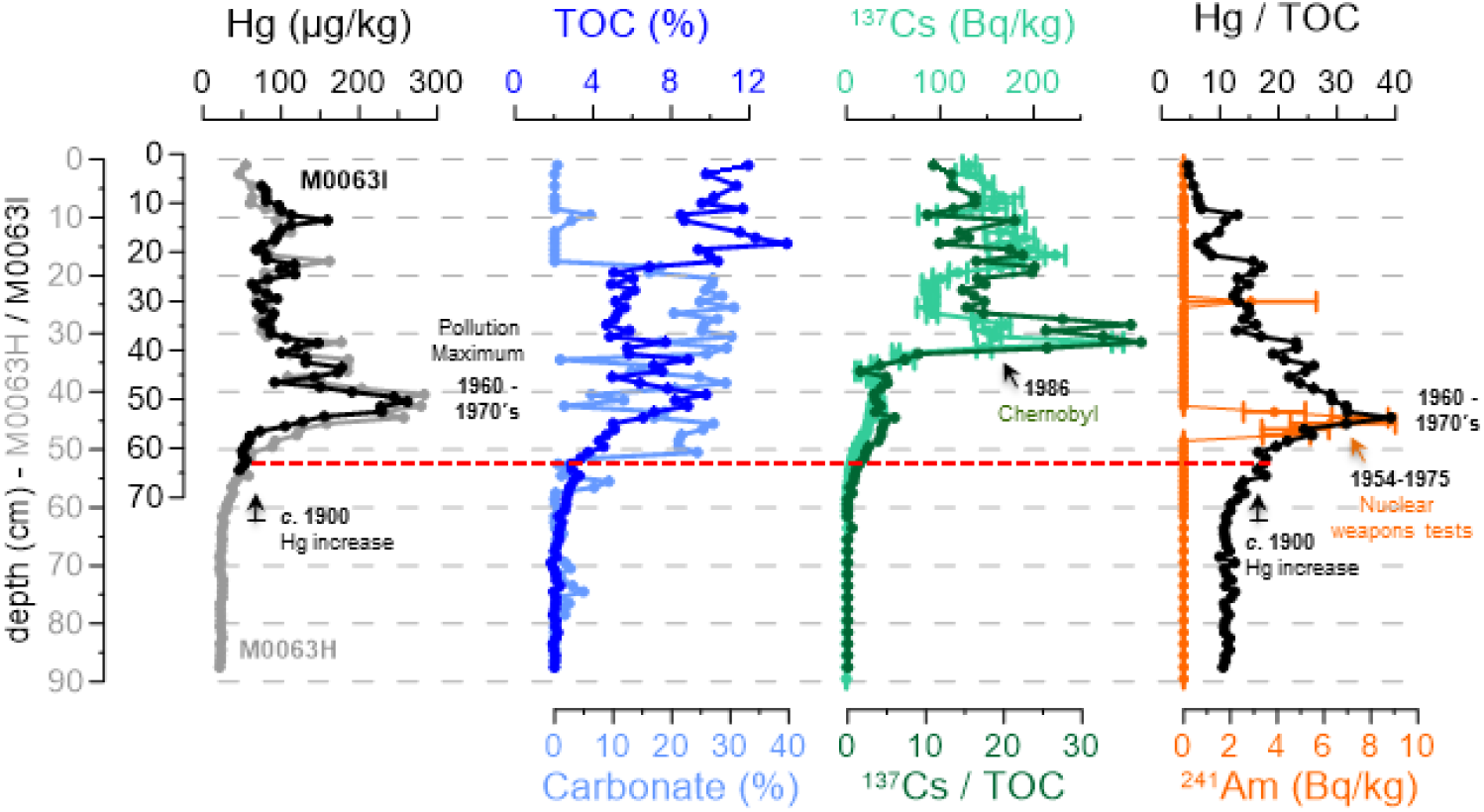
Stratigraphic time markers in cores M0063I (mercury - Hg) and M0063H (Hg and artificial radionuclides ^137^Cs and ^241^Am) following the approach described in Moros et al. (2017). Core correlation is based on Hg downcore profiles in M0063I (black) and M0063H (grey) which are shown versus respective core depths. In addition, total organic carbon (TOC, blue) and carbonate (light blue) data of M0063H are shown. Hg (dark green) and ^137^Cs (black, right) are normalized to TOC as the manganese-carbonate layers dilute the signals. Horizontal dashed red line marks an age of c. AD 1940 which can be assigned to this depth.

**Table 1.** AMS radiocarbon dates and calibrated ages of bulk sediment samples from Expedition 347, site M0063, hole C and D.

| Laboratory ID | Site and hole | Sample ID IODP | Depth (mcd) | <sup>14</sup>C Age | Error | Calibrated age 2 sigma range |  |  |
| --- | --- | --- | --- | --- | --- | --- | --- | --- |
|  |  |  |  |  |  | median | min | max |
| Core top |  |  | 0 |  |  | -63 |  |  |
| Beta 418207 | M0063C | 347-25772 | 5.22 | 1760 | 30 | 962 | 577 | 1373 |
| Beta 418209 | M0063C | 347-25774 | 6.22 | 1900 | 30 | 1301 | 872 | 1768 |
| Beta 418041 | M0063D | 347-28991 | 15.65 | 4590 | 30 | 4385 | 3690 | 4966 |
| Beta 418042 | M0063D | 347-29274 | 17.29 | 5120 | 30 | 5077 | 4533 | 5578 |
| Beta 418043 | M0063D | 347-29319 | 20.13 | 5480 | 30 | 5612 | 5066 | 6154 |
| Beta 418046 | M0063D | 347-30771 | 27.45 | 7380 | 30 | 7604 | 7040 | 8099 |

**Table 2.** Calibrated ages modelled for the diatom samples analyzed of MCD and HTM age.

| Site and hole | Sample ID IODP | Depth mcd | Calibrated age 2 sigma range |  |  |
| --- | --- | --- | --- | --- | --- |
|  |  |  | median | min | max |
| M0063C | 347-39293 | 5.42 | 1030 | 693 | 1416 |
| M0063C | 347-39299 | 6.02 | 1233 | 854 | 1641 |
| M0063C | 347-39300 | 6.12 | 1267 | 864 | 1703 |
| M0063C | 347-39301 | 6.22 | 1301 | 872 | 1768 |
| M0063C | 347-39302 | 6.32 | 1335 | 880 | 1833 |
| M0063D | 347-29645 | 21.63 | 6022 | 5579 | 6459 |
| M0063D | 347-29748 | 21.83 | 6076 | 5653 | 6499 |
| M0063D | 347-29750 | 23.74 | 6595 | 6234 | 6950 |
| M0063D | 347-30184 | 24.07 | 6685 | 6321 | 7035 |
| M0063D | 347-30204 | 26.05 | 7223 | 6760 | 7640 |

### Resting spore concentration across evolutionary timescales

We reconstructed the *Chaetoceros* spp. resting spore concentrations and observed that the highest concentration (∼203 million resting spores per gram dry weight of sediment at 26 mcd) in the sediments deposited during the HTM, after which concentrations decreased gradually with distinct peaks during MCA and present time (Fig. 4). Preservation of diatom valves were very poor between ∼21 and 8.5 mcd, however the heavily silicified resting spores were still found in some of the sediment samples in this section, but concentrations were relatively low. Due to possible dissolution, reconstructed concentrations might not be accurate in this section. During the MCA age, resting spore concentrations increased to ca. 185 million at ca. 5.5 mcd, following which concentrations decreased again. The most recently deposited sediments revealed a pronounced increase in *Chaetoceros* sp. resting spore concentration (∼146 million/gdw). The high spore concentration in the recently deposited sediments is only comparable to the concentrations found during the HTM and MCA in our record.

**Fig. 4.**
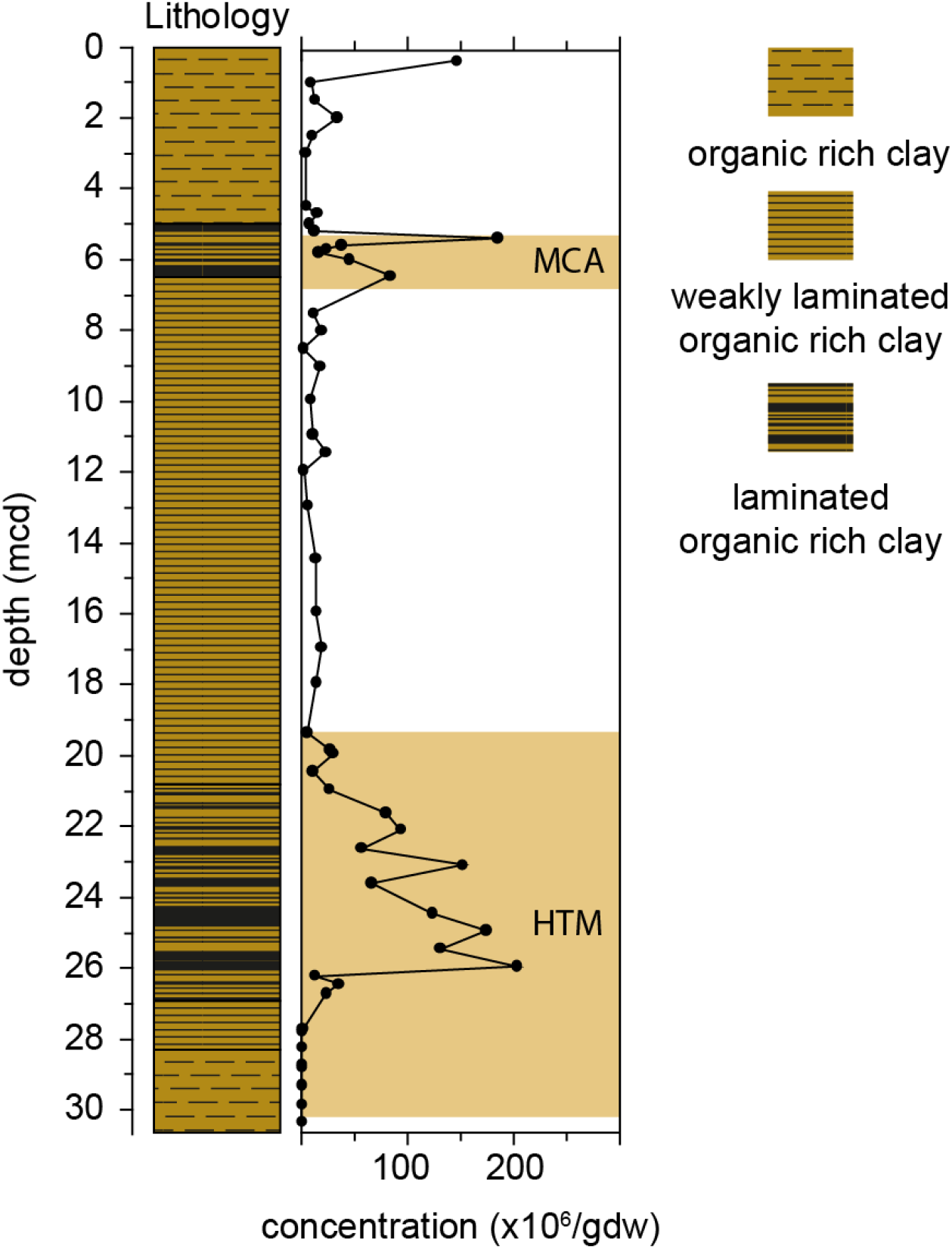
Lithology and *Chaetoceros* spp. resting spore concentrations (million valves per gram dry sediment) in the upper 30 meter sediments of IODP Exp. 347 hole M0063D (Landsort Deep). The high concentrations of MCA and HTM ages are recorded in laminated organic rich clays (equals to hypoxic bottom water conditions).

### Germination of recent and millennia old resting spores

We revived resting spores from the three time periods; (0-80 years (last century), 1300-1000 cal yr BP (MCA) and ∼7200-6000 cal yr BP (HTM). The environmental conditions were recreated to germinate the resting spores from the three ages which included reconstructing the salinity and temperature conditions. The germination of *Chaetoceros muelleri* was observed to be consistent and dominant at temperatures ranging from 15 – 18°C and at a salinity of (∼10.5-11) (Fig. 5A). On an average the time taken for the resting spores to germinate ranged from three hours to 2 −3 days.

**Fig. 5.**
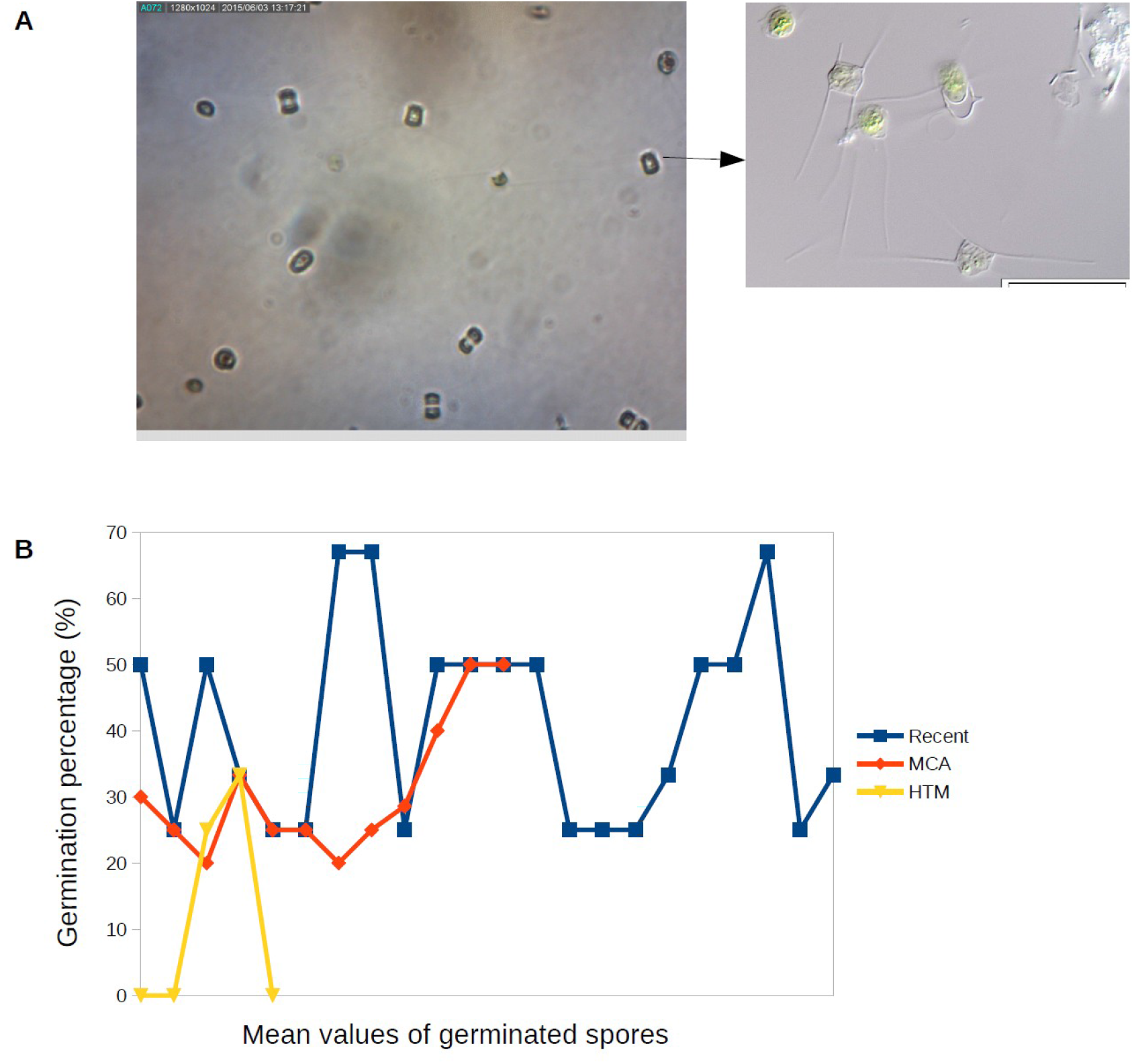
(A) Germination of resting spores of *Chaetoceros* sp. (B) Germination rate of the resting spores from the three ages.

### Germination rate and viability of recent and millenia old resting spores

The number of resting spores which germinated from the three time periods (0-80 years (last century), 1300-1000 cal yr BP (MCA) and ∼7200-6000 cal yr BP (HTM) varied. A greater number of the younger (∼0-80 calendar years and 1300-1000 cal. yr BP) resting spores germinated as compared to the oldest (∼7200-6000 cal yr BP) resting spores. On an average, the percent germination rate of the recent resting spores were the highest at 41 ± 8.7 (mean ± SE, n = 22) while the germination rate of the resting spores of the MCA and HTM ages were 31± 8.9 (mean ± SE, n= 12) and 12 ± 6.7 (mean ± SE, n = 5), respectively (Fig. 5B). A significant difference in the germination rate of the recent and ancient spores (*P* = 0.0006, F = 9.22) was found.

### Physiology and reproduction of the recent and millenia old resting spores

The resting spores isolated from the last century (0 – 80 years) reproduced and could be grown in culture but the old spores (∼1300-1000 and ∼7200-6000 cal yr BP) of the MCA and HTM age germinated but could not be grown in cultures. The time taken for the recent and ancient spores were the same ranging from three hours to 2 to 3 days, but the germination rate (%) of the ancient spores decreased with increasing time (see section on germination rate) (Fig. 5B).

### Identification of the Chaetoceros species

We identified the germinated Chaetoceros resting spores that we were able to culture as Chaetoceros muelleri Lemmermann (Fig. 6). Chaetoceros muelleri unlike other Chaetoceros taxa is a solitary living species with their vegetative cells characterized by a convex or flat valve surface (Fig. 2A-H), and their resting spores smooth with one valve bowed and the other protruded and truncated (Johansen and Rushforth1985). The above characteristics also fit with the resting spores found in the culture (Fig. 2 A-H). The taxa has been reported from brackish waters in Europe and North America (Johansen and Rushforth 1985; Guiry 2017). In the checklist of Baltic Sea Phytoplanktic species; C. muelleri is recorded in nearly all subareas and considered a warm water species found in low salinity and mainly eutrophic waters (Hällfors 2004).

**Fig. 6.**
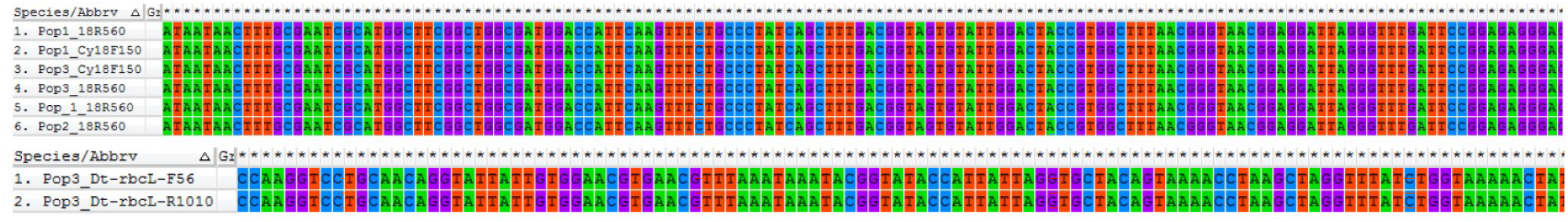
Sequences and alignment of the rbcL and SSU rDNA of *Chaetoceros muelleri*.

### Extraction and amplification of DNA from unialgal cultures of recent resting spores

We extracted high quality DNA with concentrations ranging from ∼100 to 235ng/ul with a A260/A280 ratio of ∼1.7 to 2 from the cultures of the germinated resting spores of the last century (∼0 - 80 years). We amplified six primer pair combinations from rRNA (SSU) molecules and chloroplast genes (*rbcL*) from three unialgal cultures (Tables 3).

**Table 3.**
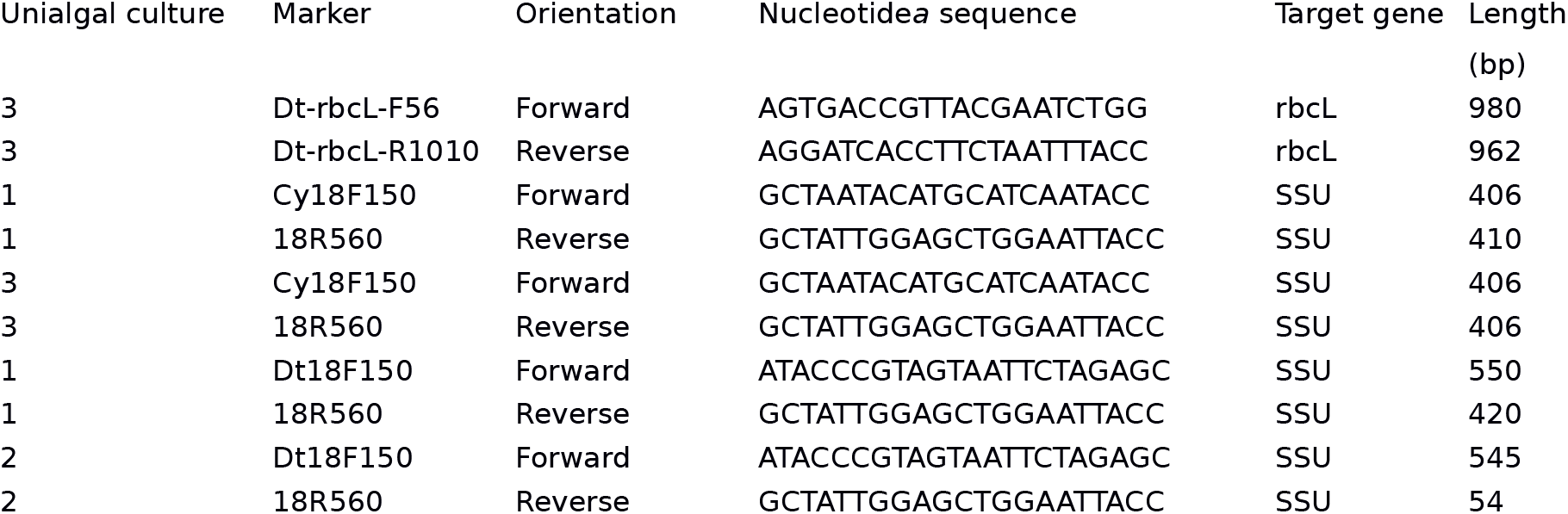
Sequence results of rRNA (SSU) molecules and chloroplast genes (rbcL) of three unialgal cultures.

### Extraction and amplification of DNA from germinated single cells of millenia old resting spores

Single cell Chelex® DNA extraction on the older spores (∼1300-1000 cal yr (MCA age)) with a A260/A280 ratio of ∼1.7 to 2 was done. DNA amplification was done using chloroplast (rbcL) and rRNA (SSU) markers.

### Sanger sequencing of DNA from unialgal cultures of recent resting spores

Partial sequences for chloroplast genes, rbcL (∼ 403- 980 bp) and rRNA molecules, SSU (406–550 bp) were obtained by Sanger sequencing for three sediment revived resting spore populations of *Chaetoceros* (Fig. 6), the sequences for the remaining genes were not obtained (Table 3). Sequence alignment and blast results of the nuclear ribosomal RNA (small subunit [SSU]) confirmed that the sequences were from the *Chaetoceros* species. Blast searches using the new sequences resulted in matches consistent with the genus-level morphological identifications of our specimens. Our validation of methods in three unilalgal cultures showed no differences in the overall sequences and alignments.

## Discussion

This study demonstrated the (i) successful germination and the germination rate of recent (∼0-80 years) and ancient resting spores (∼7200-1000 cal yrs BP) (Figs. 5A-B) (ii) DNA extraction and amplification from unialgal cultures of the recent spores (∼0-80 years) and the single cells of millennia old (∼1300-1000 cal yr BP) sediment revived resting spores (iii) DNA sequences from unialgal cultures of the recent spores (Fig. 6), and (iv) a method to identify the species (*Chaetoceros muelleri*) from the morphology of viable germinated resting spores (Fig. 2) useful in the fields of microevolutionary, micropaleontology and paleoecology studies.

By reviving resting spores deposited in sediments and extracting DNA from them we have demonstrated that *Chaetoceros muelleri* resting spores provide a promising model system for understanding species adaption and evolution across geological timescales and address questions about the impact of human-induced perturbations on life on Earth. Furthermore, previous attempts to use resting spores for paleoenvironmental reconstructions by assigning them to species met with limited success as most of the resting spores are hard to classify (Ishii et al. 2011; Witak et al. 2011), severely hindering our understanding of paleoproductivity and the effect of environmental change on species. We revived several *Chaetoceros* species but we cultured only *C. muelleri* in this study. Resting cells of diatoms are virtually identical to vegetative frustules in thickness as well as shape and pattern (McQuoid and Hobson, 1996). Previous studies on the diatom *Skeletonema* have shown that the morphology of the resting cells and germinated cells is indistinguishable (Itakura et al. 1992), whereas the resting and germinated spores of *Chaetoceros* species can be distinguished based on morphology. Therefore, the ability to identify *Chaetoceros* species from the morphology of viable germinated recent and ancient resting spores of *Chaetoceros* will improve the scope of paleoecological studies (Witak et al. 2011).

To the best of our knowledge this is the first study reviving thousands of years old (∼7200-1000 cal yr BP) diatoms from dated sediments. Previous studies on the diatom *Skeletonema marinoi* showed that the sediments were dated to be a century old (Härnström et al. 2011). Other studies on the persistence of diatoms in marine sediments have reported germination of viable resting stages from sediment layers ∼30–40 cm below the sediment surface which were estimated to be 175–275 years old (based on the sedimentation rates of 1.2–1.5 mm per year) (Stockner and Lund 1970).

Interestingly, our results indicate a different germination rate compared to the diatom species *Skeletonema marinoi.* The germination of resting spores of *Skeletonema marinoi* from sediment layers older than a few decades can take weeks to months (Härnström et al. 2011) while the recent and ancient *Chaetoceros muelleri* spores in this study germinated in three hours to two to three days, which is a surprising and promising result. However, it was observed that although the time (three hours to 2-3 days) taken for the recent and ancient resting spores to germinate remains the same the germination rate (% of germination) of the ancient spores decrease with increasing time (Fig. 5B). We also found that the older spores (∼1000-1300 cal yrs BP) could be germinated but could not be grown in cultures. It is possible that the older spores had the ability to survive but were unable to reproduce. The lower metabolic activity in the ancient spores indicates towards a trade-off between the longevity and viability of the spores and reproductive ability of the spores. The germination rates did vary significantly (*P* = 0.0006) between the different ages (Fig. 4B), which also suggests that just to remain viable over thousands of years has been a challenge for the species. The anoxic condition in the Landsort Deep possibly reduced their metabolic activity through time and helped in preserving the spores (Fig. 4). Also, it’s possible that the genetic and hence the physiological changes due to climatic or environmental changes over a thousand years could have affected its ability to reproduce.

The successful DNA extraction and amplification from unialgal cultures of the recent spores (∼0-80 years) and DNA extraction of the single cells of millennia old (∼1300-1000 cal yr BP) sediment revived resting spores reveals that the DNA in the sediment revived resting spores are well-preserved and hence presents itself as an excellent model system for tracing the evolutionary history of species. The amplification and Sanger sequencing of the plastid-encoded rbcL (∼403-980 bp) and nuclear-encoded SSU rRNA gene (406–550 bp) from the cultures show the possibilities for further studies. The non-existing variation within this genomic region was expected as this is a slow evolving region (Patwardhan et al. 2014) and was mainly used for species identification and PCR-amplification control.

Previous studies especially in humans has revealed that despite treatment of the laboratory equipment with bleach, UV irradiation of the entire facility, protective clothing and face shields and other routine precautions, contamination is a continuous threat, if for no other reason than because the specimens themselves might be contaminated with modern DNA (Hofreiter et al., 2001). Hence, the availability of high quality DNA from sediment revived recent and ancient spores in a system so well preserved where the chances of contamination are minimal as opposed to systems like humans is like a dream come true and provides immense potential for future microevolutionary studies. In this study precaution against contamination was taken in every step from sampling to PCR, by using perfluorocarbon added in the liner fluid to trace possible contamination when sampling the long cores and no contamination was detected. These cores were carefully subsampled by the curators at IODP Core Repository in Bremen and single spores were isolated with care following the protocol of Throndsen (1978). For the DNA extraction and amplification care was taken to avoid contamination; preparations were conducted in PCR-free environment.

The findings of this study reveal the potential of identifying more markers, assembling a complete genome, making comparisons of populations across temporal and spatial scales genetically with sediment revived recent and ancient spores but also experimentally with recent spores. Adequate comprehensive studies of population genetic structure in phytoplanktic species along environmental gradients like salinity combined with experimental data are lacking and the results in this study creates this opportunity. A previous study has shown that Baltic Sea populations displayed reduced genetic diversity compared to North Sea populations. The study also showed significant differences in the growth of low and high native salinity isolates indicating local salinity adaptation (Sjöqvist et al. 2015). Here we show that it will be possible to examine the genetic diversity and evolutionary changes in *Chaetoceros* populations across evolutionary timescales: ∼80 years (last century), ∼1300-1000 cal yr BP (MCA) and ∼7200-6000 cal yr BP (HTM) from sediments drilled in the Landsort Deep. Future studies can include identification of genetic markers and differences in gene expression levels to determine the changes in the population genetic structure across environmental gradients over time in sediment revived *Chaetoceros* populations. Population genomics studies using DNA data obtained from modern DNA technologies and single cell genomics together can be used to understand the evolutionary changes in the species due to natural environmental change and/or climate change or anthropogenic effects and to gain insights into the mechanisms of molecular evolution.

In conclusion, this study presents a new model system using *Chaetoceros muelleri* to address the important topic of adaptive evolution in marine species due to climate and environmental change induced by anthropogenic perturbations across evolutionary timescales. Our study reports (1) the revival, viability and germination rate of recent (0-80 years) to ancient (∼7200 - ∼1000 calendar years BP old) resting spores of *Chaetoceros muelleri* (2) the extraction of DNA and amplification of chloroplast and ribosomal genes from recent spores and DNA extraction from ancient resting spores of *C. muelleri* (3) Radiocarbon dating along with stratigraphic time markers to determine age of sediments (4) Identification of the species by reviving the resting spores (5) Baltic Sea an excellent ecosystem to study long-term effects of environment on species adaptation.

## Acknowledgement

We thank the captain and crew of R/V *Greatship Manisha* and M/S *Fyrbyggaren*, as well as the IODP Expedition 347 science party. Part of the sediment samples for this study were provided by IODP. We also thank Dr. Lubna Elabbas for assistance in the lab and Asst. Prof. Sunčica Bosak, University of Zagreb for help with species identification of *Chaetoceros muelleri*. Prof. Martin Jakobsson at Stockholm University generously made the Baltic Sea map available for our use.

